# FibrilPaint to determine the length of Tau amyloids in fluids

**DOI:** 10.1101/2023.10.13.562220

**Authors:** Júlia Aragonès Pedrola, Françoise A. Dekker, Tommaso Garfagnini, Guy Mayer, Margreet B. Koopman, Menno Bergmeijer, Friedrich Förster, Jeroen J. M. Hoozemans, Henrik Jensen, Assaf Friedler, Stefan G. D. Rüdiger

## Abstract

Tau aggregation into amyloid fibrils is linked to the development of neurodegenerative diseases, including Alzheimer’s Disease. The molecular processes underlying aggregation in disease are poorly understood. Here, we introduce FibrilPaint1 as a tool to measure the size of Tau amyloid fibrils in fluids, from early aggregation stages to mature fibrils. FibrilPaint1 is a 22mer peptide with many exciting properties, which makes it a tool for diagnostics and an attractive start point for developing a class of effective fibril targeting degraders: (i) FibrilPaint1 binds fibrils with nanomolar affinity; (ii) it does also bind to oligomeric precursors, down to a size of only 4 layers; (iii) it does not bind to monomers (KD > 100 µM); (iv) it is fluorescently labelled, which allows monitoring and localising interactions. (v) FibrilPaint1 recognises various Tau fibrils, including patient derived fibrils from Alzheimer, Corticobasal degeneration and Frontotemporal dementia; (vi) FibrilPaint1 is selective for the amyloid state and does not have background binding to amorphous aggregates, blood serum or cell lysate. In combination with Flow Induced Dispersion Analysis (FIDA), a microfluidics technology, we determined the molecular size of amyloid fibrils with sub-microliter sample volumes. This set-up acts as a molecular ruler at layer resolution - we determined Tau fibril length from 4 to 1100 layers in solution. This is an interesting parameter that can be used for diagnostic applications and biochemical research in dementia.

## Introduction

The aggregation of proteins into amyloid fibrils characterizes the development of neurodegenerative diseases, that affect 55 million people worldwide ^1,2^. Amyloids are long, fibrillar structures with a typical cross-β fold ^3–5^. Amyloid formation of the protein Tau is related to the progression of several Tauopathies, including Alzheimer’s Disease (AD), Frontotemporal Dementia (FTD), and Corticobasal Degeneration (CBD) ^6–8^. Tau is an intrinsically disordered protein that can aggregate into distinct conformations when detached from the axons of neurons^4,8–11^.

The formation of amyloid fibrils is one of the earliest pathological hallmarks of neurodegenerative diseases ^3,6,12–15^. Recently, the antibodies Aducanumab and Lecanemab have been approved to target Amyloid-β (Aβ) plaques in the brain, aiming to slow down the progression of the disease and improve cognitive function ^16,17^. In AD, Aβ aggregation outside the cell precedes Tau aggregation inside the cell^4,13,18^. The effects of these antibodies remain modest, with significant risks involved, and most patients are not eligible for these therapies as they apply only to very early stages of the disease ^16,17,19,20^. Future therapies and treatments would strongly benefit if they could be administered early, preferentially before first symptoms appear ^18,21–23^.

Molecular understanding of aggregation is crucial for the development of new causal therapeutic strategies. Key for this is a quantitative understanding of the underlying nucleation and elongation phases of the process ^24,25^. Nucleation is the formation of a seed that starts the aggregation process^26,27^. Such nuclei that initiate the aggregation reaction form during a lag-phase populated by transient and highly dynamic states of the protein. Elongation is the stacking of fibril layers on top of each other, lengthening the fibrils^25,27^.

Accurate measurement of these steps remains a challenge. Small amyloid fibrils among these first nucleation steps easily escape detection, although already present in the lag-phase ^28,29^. Elongation remains challenging to characterize accurately. Amyloid dyes can estimate the presence of amyloids but they cannot provide information of the size of the amyloid fibrils^30–33^. Microscopy techniques visualise fibril structures down to atomic resolution but they are limited in scalability and quantitative readout. To reveal a quantitative picture of the aggregation process, it is desirable to develop new sensitive and quantitative microscale methods in solution.

Here we present a method to measure the length of Tau amyloid fibrils in solution. We developed FibrilPaint1, a peptide that specifically binds to Tau amyloid fibrils, but not monomers. In combination with Flow Induced Dispersion Analysis (FIDA), we determine the fibril length of these Tau fibrils in fluids, such as blood plasma. We show its applicability to follow the elongation of recombinant TauRD, the repeat domain of Tau protein, over time. We also used this set-up to determine the length of end-stage Tau fibrils from patients diagnosed with AD, FTD and CBD.

## Results

### Development of a method to detect Tau amyloids in fluids

We set out to develop a method to detect Tau amyloid fibrils (**Fig. 1**). To do so, we need a specific tracer to bind these Tau fibrils. Based on our previous work ^34^, we designed a set of four new peptides with enhanced binding to amyloid aggregates. We increased the π-stacking and H-bonding residues of the peptides^34 35^. In addition, we added a negatively charged EEVD sequence C-terminally with a GSGS spacer, creating two oppositely charged sections, with a positively charged N-terminal region and a negatively charged C-terminus. The EEVD sequence may also serve as adaptor to recruit protein quality control factors that specifically recognise this sequence via a TPR domain, which is potentially interesting for follow-up studies ^36,37^. We added a fluorescein group (Fl-) at the N-terminus to allow detection. We varied the net charge and the number of aromatics in the peptides. This further optimization resulted in a set of four 22-residue long peptides with different charge, number of aromatics and distribution of the different types of residues (**Table 1**). Designed to make fibrils visible, we call this class of peptides FibrilPaint. FibrilPaint1 and FibrilPaint2 have Arginine residues on positions 4 and 5, whereas FibrilPaint3 and FibrilPaint4 have Aspartates. FibrilPaint1 and FibrilPaint4 have Histidines on positions 11 and 12 where FibrilPaint2 and FibrilPaint3 have Threonines.

**Figure 1.**
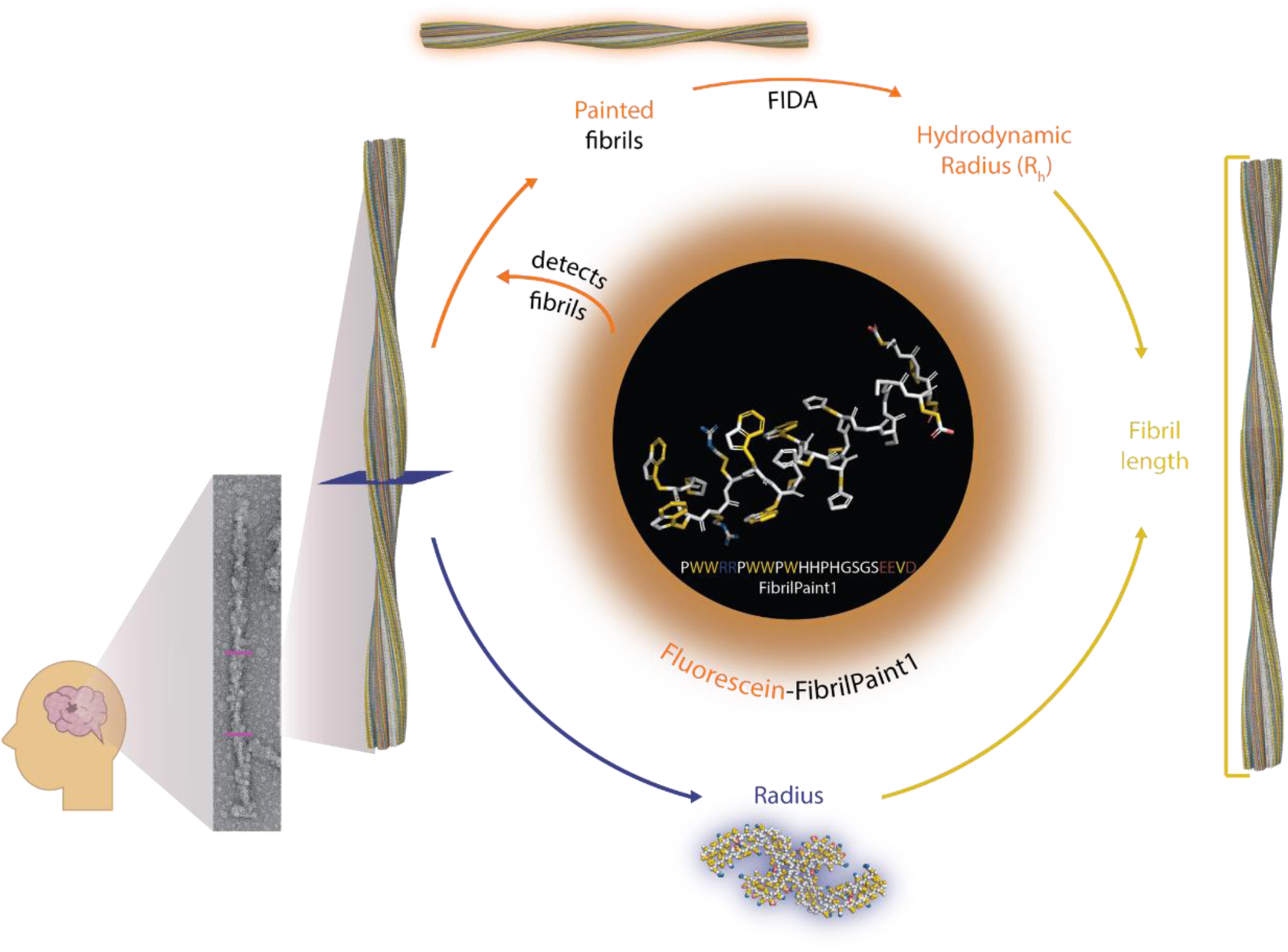
Abstract Figure: FibrilPaint1 to detect the presence of fibrils and determine fibril size. FibrilPaint1 (in circle) detects the presence of amyloid fibrils. Through its Fluorescein-tag, binding works as fluorescent staining of fibrils (orange pathway). The stained fibrils can be analysed with FIDA to get the hydrodynamic radius (R_h_) of the fibril population. If the structure of the fibril is known, this can be used to determine the radius of the fibril (blue pathway). Otherwise, estimations of the radius can be made. Combined, these data can be converted to give the fibril length (yellow pathway).

**Table 1.**
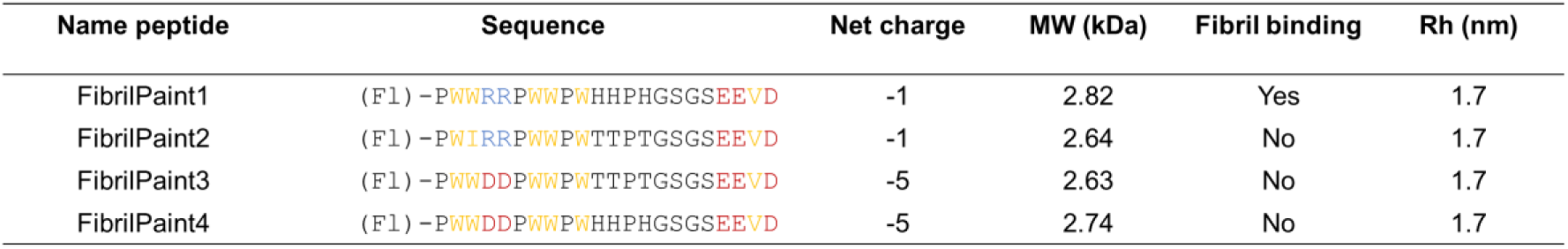
Peptide names and sequences. (Fl) at the N-terminal of the sequence indicates fluorescein, which is not considered when calculating the molecular weight or number of amino acids. Amino acid colouring is based on YRB colours (hydrophobic residues, yellow; negatively charged residues, red; positively charged residues, blue) ^38^ Net charge is the total charge of the different peptides at physiological pH. Fibril binding is the detection of Tau amyloid fibrils with Flow Induced Dispersion Analysis (FIDA). Hydrodynamic Radius (R_h_) is the the radius determined with FIDA when the peptides do not bind anything.

### The four peptides interact with Tau fibrils

We used Tau Repeat Domain (Q244-E372, TauRD) with the pro-aggregation mutation ΔK280 as a model for screening our FibrilPaint compounds. We performed a Thioflavin T (ThT) assay. ThT is an amyloid dye and provides a semi qualitative methods used for the identification of amyloid structures *in vitro* and histology staining ^32,33,39^. A ThT assay is an established method to monitor protein fibrils formation, in which the fluorescent dye ThT emits fluorescence upon binding to fibrils ^39^. The four FibrilPaint peptides decreased the emitted ThT signal significantly, at substoichiometric concentration in a dose-dependent manner (**Fig. 2A**). This demonstrates the FibrilPaints interact with the TauRD fibrils.

**Figure 2.**
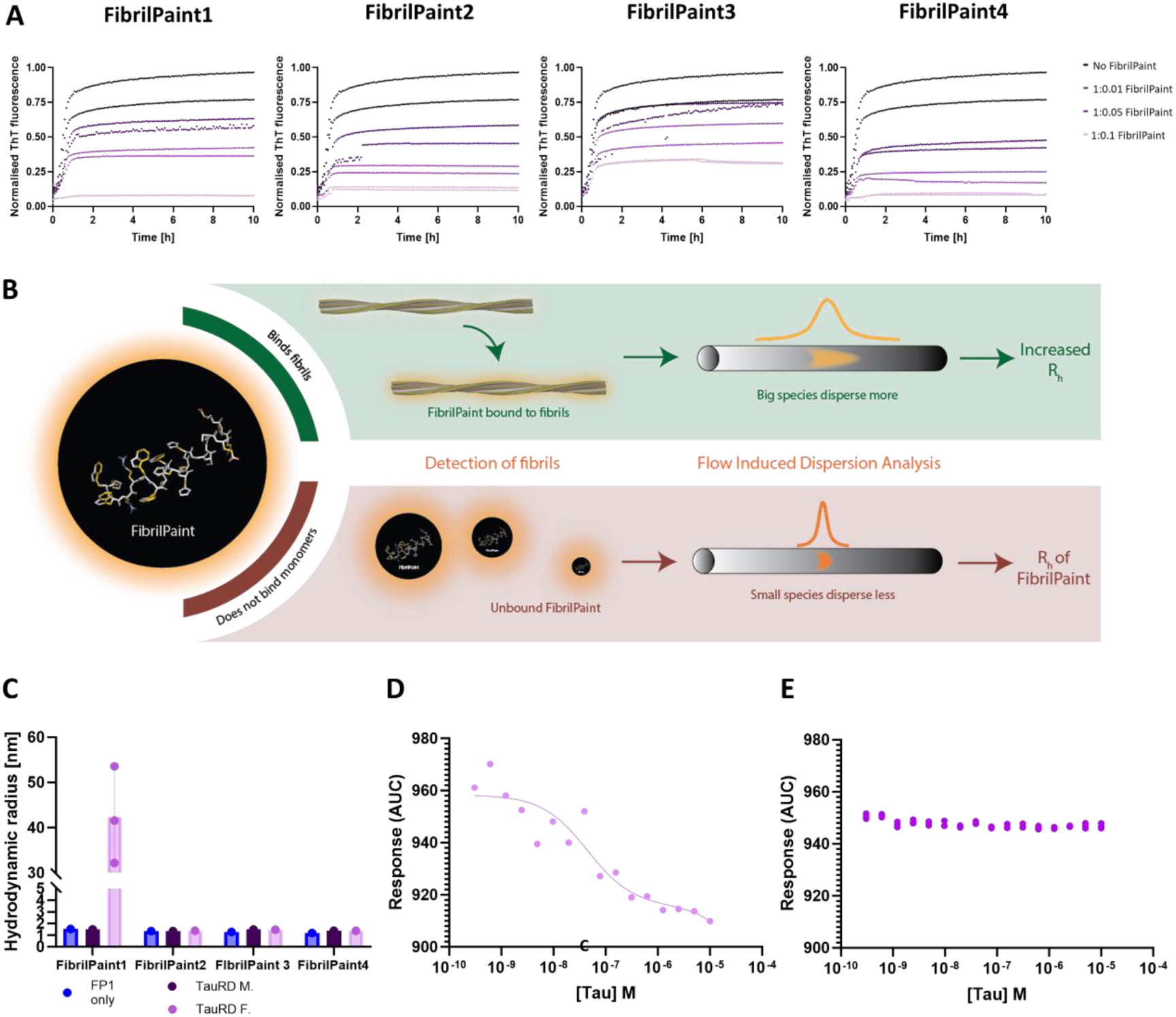
FibrilPaint1 is a specific label for protein fibrils. **A)** Representative figure of ThT assays with 20 μM TauRD aggregation and titration of 2 - 0.2 - 0.02 µM FibrilPaint1, FibrilPaint2, FibrilPaint3 or FibrilPaint4 (left to right). All peptides lower the end-plateau in a dose-dependent manner.Triplicates are shown. N=3 **B)** Schematic of FibrilPaint binding to fibrils with Flow Induced Dispersion Analysis (FIDA). If a FibrilPaint binds, this leads to a bigger species, which disperses more in FIDA, leading to an increase in the perceived R_h_ (green box). If there is no binding, the R_h_ remains the same as the R_h_ of FibrilPaint only (red box). **C)** Results FIDA of FibrilPaint binding to monomeric or fibrillar TauRD. Average of N=3 shown **D)** Binding curve of FibrilPaint1 with titrated TauRD fibrils, or **E)** monomers N=3. (FibrilPaint peptide only, blue; TauRD monomers, dark purple; TauRD fibrils, light purple;).

### FibrilPaint1 binds to Tau fibrils

Next, we screened the four FibrilPaint peptides for direct binding to TauRD fibrils. We developed an application of FIDA to determine the size of amyloid aggregates (**Fig. 2B**). FIDA is a recently established technique to measure size of protein complexes^40,41^. In FIDA, the detector records the fluorescence signal after passage of the sample through a long capillary. The sample passes a detector window, resulting in a size-dependent diffusion profile of the fluorescent compound. Smaller species diffuse faster and bigger ones slower, resulting in a narrow or broad dispersion of the fluorescent signal (**Fig. 2B**). Since FIDA readout relies on the physical diffusion properties of the sample, we can measure the hydrodynamic radius (R_h_) of labelled species from the fluorescent signal^42–44^. The R_h_ is a parameter corresponding to the size and shape of a molecule or a complex in solution. It is the radius of the sphere created by the tumbling particle. R_h_ values can be estimated from the structural coordinates, either by experimental methods or by predictions such as AlphaFold (**Table 2**). For known structures, e.g. for many protein fibrils, an experimentally determined R_h_ value allows to estimate the dimensions of the particle.

**Table 2.**
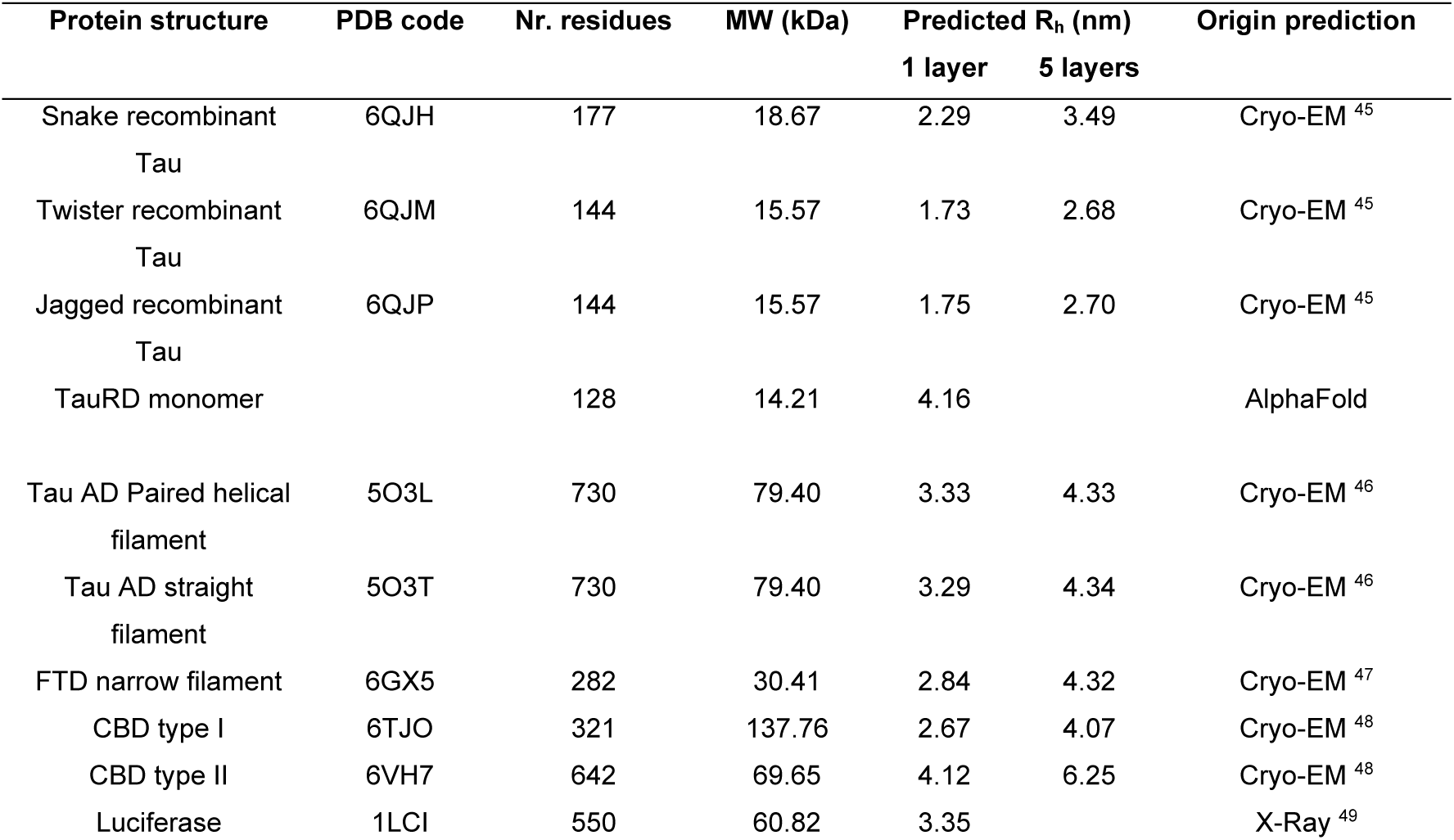
Protein structures and their predicted hydrodynamic radii. The number of residues and the molecular weight are given for the total structure as available in the PDB. If structures are available, the included 1 layer R_h_ is the R_h_ a structure would have if it could fold in its fibrillar conformation with just one layer. The 5 layers R_h_ is the R_h_ that results from a fibril that stacks 5 layers. Origin of the prediction is the method used to determine the coordinates of PDB structures.

With this set-up, we determined the R_h_ of FibrilPaint1 to 4 to be 1.7 nm each (**Table 1**). The binding of the fluorescently labelled FibrilPaint peptide to protein fibrils increases the R_h_ value, as the peptide now tumbles together with the larger fibril (**Fig. 2B**). To measure the R_h_ of fibril complexes, we incubated pre-formed fibrils together with each of the four peptides in a 1:100 peptide:monomer ratio, using 200 nM peptide with fibrils made of 20 uM TauRD monomers.

Of all the FibrilPaint peptides, only FibrilPaint1 bound to TauRD fibrils, increasing the average R_h_ value from 1.7 nm to 45 nm (**Fig. 2C**). Importantly, incubation of FibrilPaint1 with TauRD monomer did not result in an increased R_h_ value, indicating that it does bind specifically to the fibril but not to the monomer (**Fig. 2C**). It is interesting that only FibrilPaint1 binds strongly enough to act as a non-covalent label, given that all FibrilPaint peptides interact with TauRD fibrils (**Fig. 2A**). This indicates that high affinity for amyloid fibrils exceeds the demands required for ThT competition or modulation of aggregation.

We confirmed that FibrilPaint1 binds specifically TauRD fibrils and not monomers by assessing binding affinity with monolith spectral shift analysis (**Fig 2D, E**). FibrilPaint1 shows no binding affinity for monomers (**Fig 2E**), but does for TauRD amyloid fibrils (**Fig 2D**). K_d_ is estimated to be around 57 nM, based on monomer concentration. Since multiple monomers are needed to form one fibril, true affinity lies higher.

### FibrilPaint1 does not bind to amorphous aggregates

Having confirmed the ability of FibrilPaint1 to bind Tau amyloid fibrils and not monomers, we investigated whether binding of FibrilPaint1 to aggregates requires amyloid structures. We used Luciferase as an established paradigm for non-amyloid aggregates ^50^. Luciferase is a globular 61 kDa protein with a R_h_ of 3.4 nm (**Table 2**) that forms amorphous aggregates when denatured by heat shock. We incubated FibrilPaint1 together with heat-shocked Luciferase in a 1:100 ratio. The R_h_ of FibrilPaint1 in the FIDA measurement is not affected by the presence of luciferase aggregates (**Figure 3A**). This indicates that FibrilPaint1 does not bind to amorphous luciferase aggregates, demonstrating specificity for amyloid fibrils.

**Figure 3.**
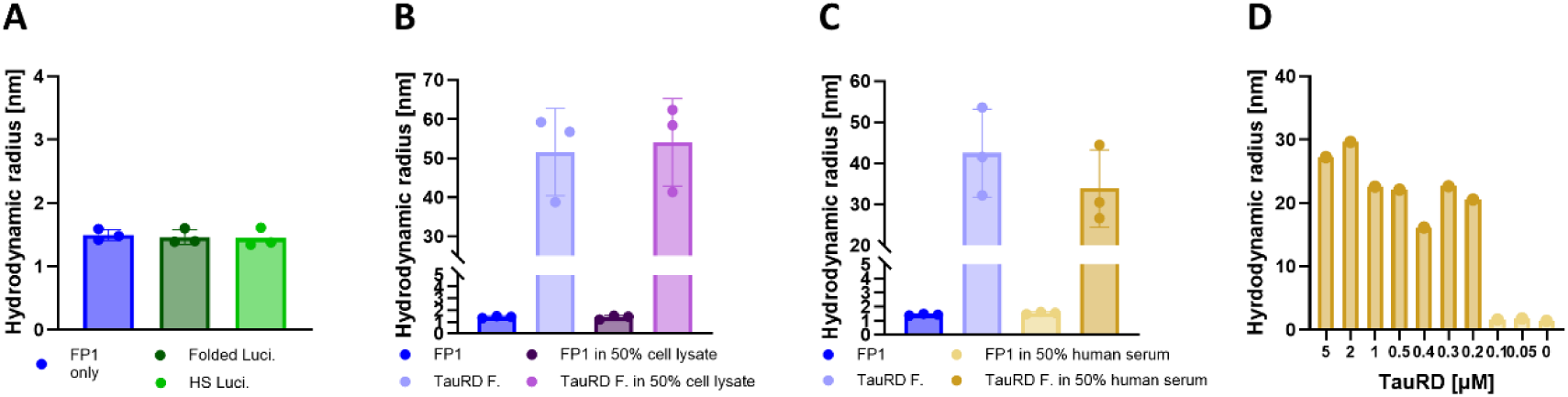
FibrilPaint1 is specific for amyloid aggregates. **A)** 0.2 μM FibrilPaint1 incubated with amorphous aggregated Luciferase by heat shock. FibrilPaint1 has a R_h_ of 1.7 nm, incubated with heat shock luciferase the size remains the same. **B)** Binding of 0.2 μM FibrilPaint1 to TauRD fibrils in buffer or 50% cell lysate. In buffer, FibrilPaint1 has a R_h_ of 1.7 nm, incubated with TauRD fibrils has an average R_h_ of 52 nm. In 50% cell lysate, FibrilPaint1 has a R_h_ of 1.7 nm, incubated with TauRD fibrils has an average size of 54 nm. **C)** Binding of 0.2 μM FibrilPaint1 to TauRD fibrils in buffer or 50% human serum. In the presence of buffer, FibrilPaint1 has a R_h_ of 1.6 nm, incubated with TauRD fibrils has an average R_h_ of 42 nm. In 50% human serum, FibrilPaint1 has a R_h_ of 1.5 nm, incubated with TauRD, it has an average size of 34 nm. (**D)** Binding of 0.2 μM FibrilPaint1 to different concentrations of TauRD fibrils in 50% human serum. The detection limit of FibrilPaint1 observed under these conditions is 200 nM TauRD monomers, corresponding to fibrils formed at 200 nM (FibrilPaint1 in buffer, dark blue; folded luciferase, dark green; heat-socked luciferase; light green; TauRD fibrils in buffer, light blue; FibrilPaint1 in cell lysate, dark purple; TauRD fibrils in cell lysate, light purple; FibrilPaint1 in human serum, light yellow; TauRD fibrils in human plasma, dark yellow).

Next, we investigated whether FibrilPaint1 binds fibrils without perturbance by other biomolecules or cellular components. We tested selectivity for TauRD fibrils in presence of an abundance of other cellular proteins, by adding *E. coli* cell lysate. We incubated pre-formed TauRD fibrils with FibrilPaint1 in a 1:100 ratio in 50% cell lysate. Under these conditions, FibrilPaint1 shows an increased R_h_ of 54 nm, which is consistent with the R_h_ of 52 nm measured from the same fibrils in buffer (**Fig. 3B**). When incubating FibrilPaint1 alone in cell lysate, the R_h_ remains the same as FibrilPaint1 alone (**Fig. 3B**). Thus, FibrilPaint1 specifically recognises fibrils in a complex cellular mixture.

To test FibrilPaint1 in a more physiological and disease-relevant environment, we incubated FibrilPaint1 in absence or presence of a new batch of pre-formed TauRD fibrils in 50% human serum. This batch has an R_h_ of 42 nm in buffer. FibrilPaint1 alone in 50% human serum appeared to be 1.5 nm, which correlates to the parallel measurement of 1.4 nm observed in buffer (**Fig. 3C**). In presence of pre-formed TauRD fibrils, the R_h_ increases up to 34 nm in 50% human serum (**Fig. 3C**). Differences in the R_h_ are most likely due to fibril heterogeneity.

After confirming the ability of FibrilPaint1 to detect TauRD fibrils in human serum, we aimed to estimate the concentration threshold for such measurements. We titrated recombinant TauRD fibrils into serum and measured the R_h_ with FibrilPaint1/FIDA for decreasing concentrations (**Fig. 3D**). TauRD fibrils in this experiment had an average R_h_ value of 21 nm, but this size varies due to the heterogeneity of amyloid fibrils. We can detect these fibrils down to a lower limit of fibrils made from a monomeric solution of 200 nM. With the measured R_h_ value of 21 nm, we can calculate the number of monomers in the average fibril. An R_h_ of 21 nm corresponds to 260 layers for a PHF-shaped fibril (**Table 2**, **Fig. 6F**). This means the true fibril concentration is as low as 400 pM. These findings indicate that FibrilPaint1 might be useful to screen for the presence of subnanomolar amyloid concentrations found in pathological settings.

### Monitoring fibril kinetics

Diagnostic applications strongly benefit from detection of early aggregating species. To evaluate FibrilPaint1’s potential as a rapid diagnostic tool, we tested its ability to detect fibrillar species formed during the early stages of aggregation. We aimed to track fibril size throughout the aggregation process by observing the progressive increase in R_h_, using FibrilPaint1 and FIDA. To do so, we performed aggregation reactions in absence of FibrilPaint1. At the desired time, we added FibrilPaint1 to the reaction.

After 0.5 h, the aggregation reaction of TauRD resulted in an increase of the averaged R_h_ from 1.7 nm, which corresponds to FibrilPaint1 alone, to 2 nm (**Fig 4A**). TauRD aggregation continued, and rapidly increased to an R_h_ of 5 nm after 2 h (**Fig. 4A**). Only the largest species were plotted (**Fig. 4A**). TauRD aggregation reached a plateau at an R_h_ value of 45 nm after 8 h.

**Figure 4.**
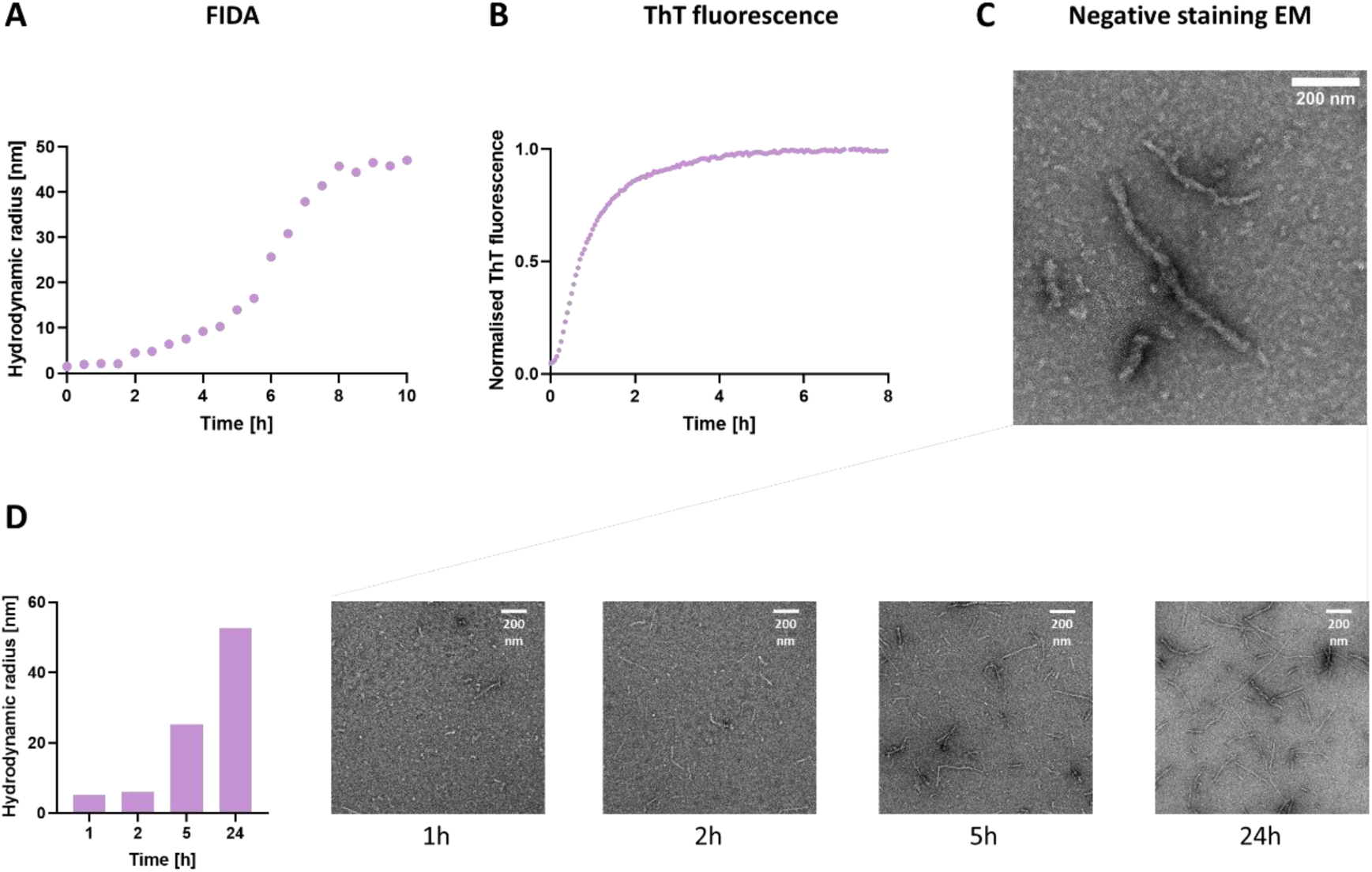
Aggregation of TauRD over time. **A)** Hydrodynamic radius of TauRD amyloid species as determined with FIDA. Representative figure shown of N=3 **B)** In parallel, aggregation was monitored with a ThT assay. Average of triplicate measurement shown, N=3 **C)** The end-product of TauRD was imaged with TEM after 24 h **D)** In parallel experiments of TauRD aggregation monitored with FIDA (left) or TEM at 1h, 2h, 5h and 24h. Average data shown of duplicate measurements.

We monitored the aggregation process in parallel with the established ThT assay (**Fig. 4B**). The ThT fluorescent signal rised immediately after addition of heparin to TauRD and increases rapidly, reaching the plateau-phase after 4 h (**Fig. 4B**). No changes were monitored with ThT anymore, but after this point, the TauRD fibrils continue elongate from an R_h_ of 10 nm to an R_h_ of 45 nm. The faster saturation of the ThT fluorescent signal suggests an equilibrium in binding and release of ThT to fibrils, while the fibrils are still growing in length. The FIDA measurements provide an interesting additional readout which can monitor the changes in aggregation for a longer period.

### Correlating hydrodynamic radius to fibril length

Next, we compared the average fibril size obtained by FIDA data with TEM images. We used TEM imaging to characterise the shape of the TauRD fibrils after 24 h aggregation (**Fig. 4C**). They appeared as long, single, fibrillar structures (**Fig. 4C**). We also monitored TauRD fibril formation using transmission electron microscopy (TEM) (**Fig. 4D**). After 1 and 2 h of aggregation, only a minimal number of fibrils were observed, due to the resolution of the technique, which limits the detection of smaller species. After 5 h, more and longer fibrils became visible, which elongated further up to 24 h.

To correlate the readout of TEM to our FIDA measurements, we built a model to convert the hydrodynamic radius to fibril length. For Tau, many different cryo-EM structures are now available, providing us conformational information^46,47,51,52^. For TauRD however, there is no cryo-EM structure available. We chose to use the PHF of AD as a model. The length of the PHF can be determined through its turns, which we used to verify our model. One full turn of the twist of a PHF comprises 342 layers. As the fibril structure is per layer 0.47 nm, full twist corresponds to a length of approximately 160 nm ^46^. We stacked *in silico* layers on top each other and calculated the predicted with the R_h_ with FIDAbio R_h_ prediction tool (**Fig. 5A**). As a result, we have a mathematical equation to convert the R_h_ to fibril layers and length (**Fig 5B**).

**Figure 5.**
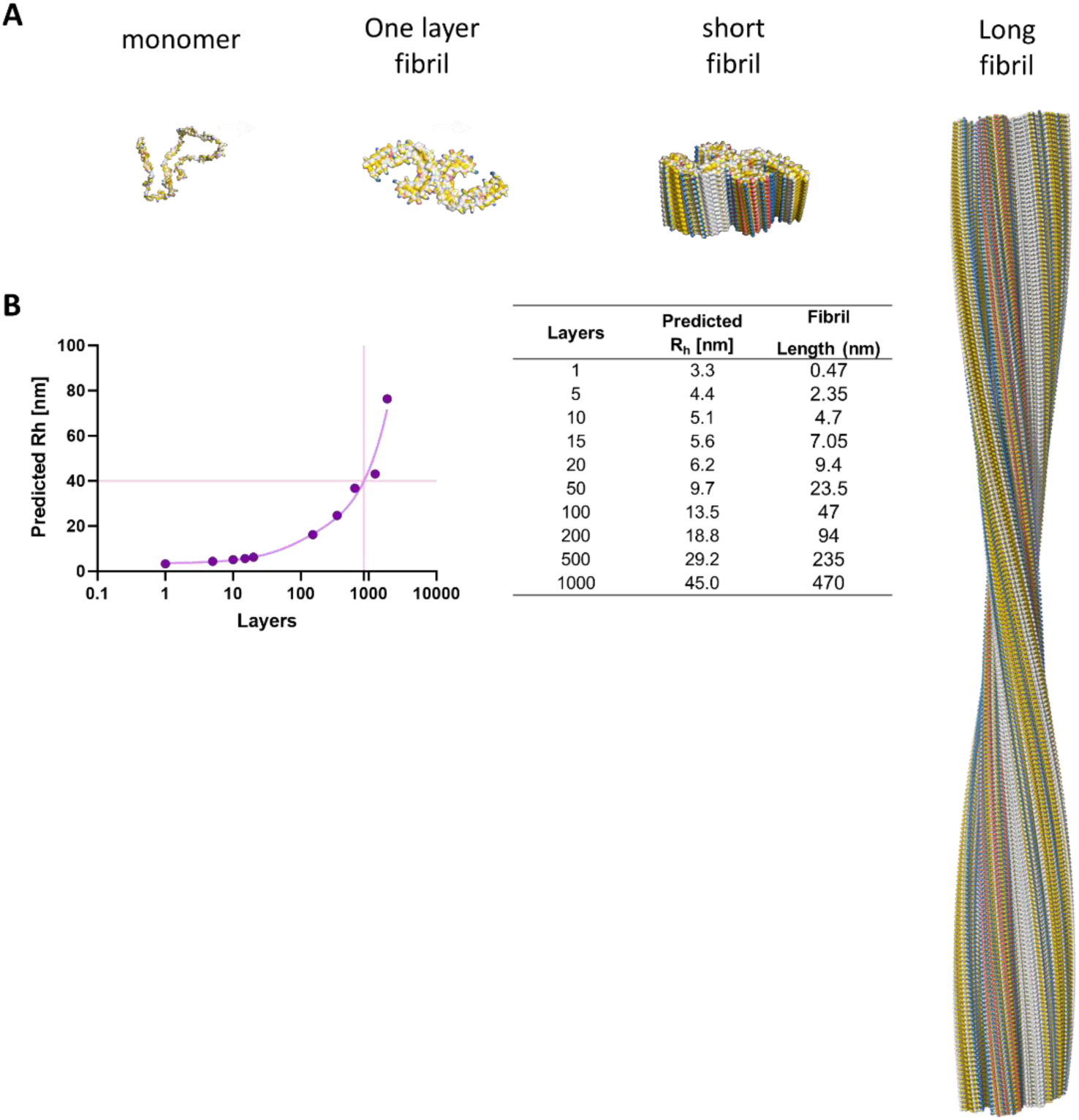
Correlation between fibril length and hydrodynamic radius (R_h_). **A)** Structures of Tau monomer and Paired Helical Filament fibril as one layer, 20 layers and 342 layers. **B)** R_h_ prediction based on the coordinates of the stacked fibril layers. Fibril length is calculated using 0.47 nm for one layer ^46^.

The length of TauRD fibrils at 24h varied, 500 to 600 nm on average (**Fig 4C**). This is slightly longer than we find with FIDA, where we measure a length of roughly 470 nm (**Fig 4A**). This is expected, as smaller fibrils remain undetectable due to the limited sensitivity of TEM for smaller particles.

The need for measuring fibrils directly in fluid becomes clear at the start of the aggregation process. The resolution of TEM limits the visibility of fibrils below 100 nm. At the earliest timepoints, almost no fibrils can be found with that length. In our microfluidics set-up, all fibrils were measured, also the short ones. At 1h, an average R_h_ of 4 nm was determined. Assuming a PHF fibril structure, this correlates to approximately 4 layers, and a fibril length of 2 nm (**Fig 5B**). After 2 h, the R_h_ increased 5 nm, corresponding to approximately 10 layers and a fibril length of 4.7 nm (**Fig 5B**). It is however likely that early fibrils take on other conformations with smaller interfaces ^28,29^. Thus, the FibrilPaint1/FIDA assay allowed detecting early species involved in aggregation, offering the possibility to screen for Tau amyloid fibrils early in the disease progression.

### FibrilPaint1 detects patient-derived fibrils

As potential reporter for Tau fibrils, it is important to assess the ability of FibrilPaint1 to recognise patient-derived fibrils of several Tauopathies. Interestingly, cryo-EM structures of Tau fibrils show that their shape differs for the various tauopathies, and heparin-induced recombinant fibrils ^10^ (**Fig. 6B**). Therefore, we set out to measure *ex vivo* Tau fibrils from three different Tauopathies. We purified fibrils from deceased patients diagnosed with CBD, FTD and AD ^46,47,51^. Monomeric Tau undergoes different post-translational alternative splicing in different diseases, leading to six different isoforms, with either four (4R) or three (3R) repeats of the microtubule-binding domain. Depending on the isoform incorporated in the fibril, tauopathies can be classified as 4R (CBD), 3R (FTD) or 4R/3R (AD). These different isoforms also have different conformations in each disease ^10^. To confirm the typical disease-specific fibril shape, we imaged the purified fibrils with TEM (**Fig. 6C**).

**Figure 6.**
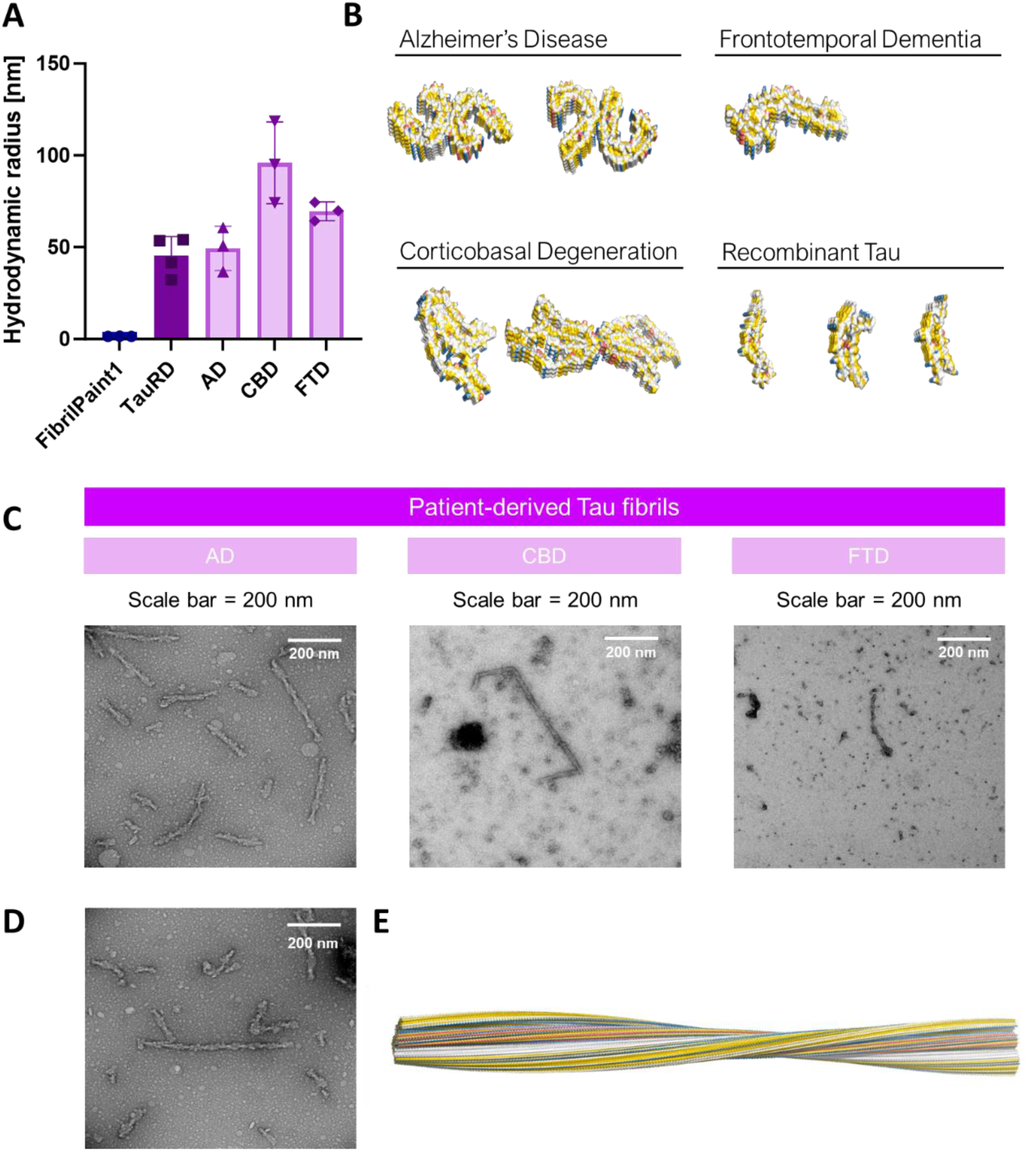
Size assessment of patient-derived material from various tauopathies. (A) Hydrodynamic radius of fibrils extracted from patients diagnosed with Alzheimer’s Diseases (AD), Frontotemporal Dementia (FTD) and Corticobasal Degeneration (CBD), measured with FibrilPaint1. (FibrilPaint1 only, blue; TauRD fibrils, dark purple; fibrils retrieved from patients, light purple). (B) Structural representation of the Tau fibrils of different tauopathies, and recombinant Tau fibrils for comparison. The images were generated in PyMol software based on the coordinates of the cryo-EM structures. Functional groups are coloured according the YRB script. PDB codes: AD paired helical filament, 5O3L; AD straight filament, 5O3T; FTD narrow filament, 6GX5; CBD type I 6TJO; CBD type II 6VH7; Snake recombinant Tau, 6QJH; twister recombinant Tau, 6QJM; Jagged recombinant Tau 6QJP (Colours in YRB: yellow; negative charge, red; positive charge, blue). (C) TEM of patient-derived fibrils from AD, CBD and FTD (FLTR). (D) TEM of PHF from AD (E) Structural representation of AD paired helical filament consisting of 342 layers. The image was generated in PyMol software based on coordinates of cryo-EM structures. Functional groups are coloured according to the YRB colour code ^38^.

Using our microfluidics set-up, we could detect all three tauopathies with our screening set-up (**Fig 6A**). The fibrils formed by recombinant TauRD had an average R_h_ of 45 nm. Fibrils measured from a patient diagnosed with AD had an average R_h_ of 49 nm, with little variation between the measurements (**Fig 6A**). For CBD, the observed R_h_ was 95 nm, with more variation in between measurements, likely reflecting a heterogeneous population of fibrils in the patient’s brain (**Fig 6A**). Fibrils from FTD appeared to be a homogeneous population, with an average R_h_ of 69 nm (**Fig 6A**).

In the case of AD, the average Rh of 49 nm corresponds to a length of 510 nm or 1100 layers, as given by our FIDAbio prediction tool (**Fig. 5B**). Using TEM, we investigated 103 fibrils and observed 48 showing the typical PHF structure and an average 2.7 turns. With the known repetitive element in these fibrils, this corresponds to 430 nm (Fig. 6E). Of course, this excludes the Straight Filaments found in the same fibril population. The order of magnitude is in line with the 510 nm observed in the FIDA experiments with the FibrilPaint1-stained fibrils. Thus, we conclude that combining FibrilPaint1 and FIDA is suitable to determine the length of pathological fibrils.

## Discussion

We have developed FibrilPaint1, a novel peptide capable of recognising Tau amyloid fibrils in solution with high affinity, both recombinant and patient-derived fibrils from three distinct tauopathies. This provides a molecular ruler for Tau fibrils over the full aggregation range, measuring fibril length from 4 to 1100 layers. FibrilPaint1 is applicable in complex environments such as human serum, where we can detect fibrils in sub-nanomolar concentrations. FibrilPaint1 is, therefore, a tool to assess the length of Tau amyloid fibrils throughout the aggregation process.

The non-covalent, high-affinity fluorescent-labelling of Tau amyloid fibrils allows studying these fibrils in complex samples. However, FibrilPaint1 is characterised by a large proportion of aromatic residues (tryptophan and histidine) and also arginine. These residues are capable of engaging in π-stacking interactions, which is consistent with earlier findings that Tau fibrils aberrantly attract proteins enriched in π-stackers ^35^. The avidity effects caused by the repetitive nature of fibrils increases the importance of such low affinity contacts ^34,35^. This is consistent with our observation that Tau monomers do not interact with FibrilPaint1. This interaction chemistry is different from other amyloid fibril binders. Some dyes such as Thioflavin T bind to various fibrils by intercalating^53^. Flortaucipir and EGCG form repetitive small molecule stacks framed by the fibril fold ^19,54^. It is unlikely that FibrilPaint1 forms such stacks given that it binds under binds under conditions when fibril-forming monomer is present at 100x excess over FibrilPaint1. The precise interactions that determine the high affinity of FibrilPaint1 are still to be determined. Uncovering the molecular basis of amyloid specificity could guide the design of next-generation tools.

As FibrilPaint1 binds with high affinity to Tau amyloid fibrils but not monomers it could be of interest for developing novel warheads for Targeted Protein Degradation (TPD). TPD is based on bifunctional degraders, of which one side binds specifically to the target, the warhead, and the other recruits a protein degradation machinery ^55–58^. The C-terminal EEVD sequence of FibrilPaint1 targets the E3-ligase CHIP, which can ubiquitinate substrates for proteasomal degradation ^58,59^. FibrilPaint1 may also be functionalised with alternative tags that can guide the tracer towards degradation.

The high affinity of FibrilPaint1 to Tau allows the determination of the R_h_ of a fibril population in solution, which provides quantitative ensemble information about Tau amyloid fibrils at nanometer resolution. This is an important parameter for assessing fibrils and for monitoring their growth or shrinkage. This goes beyond information provided by ThT or antibody-based detection assays, which cannot distinguish small from large aggregates ^30,31^. Size information in solution is complementary to imaging data obtained by microscopy techniques, which are capable of distinguishing fibril morphologies but lack scalability for widespread application ^60–62^. The FibrilPaint/FIDA assay has no background signal of serum or other complex biofluids requires sub-microliter sample volumes.

Reliable determination of fibril length in solution may offer opportunities to use this parameter as a biomarker for diagnostic purposes. This could be relevant for monitoring the effect of future amyloid-targeting drug or for stratification of patient cohorts for clinical trials. We have demonstrated the applicability of FibrilPaint1 for *ex vivo* amyloid fibrils of several Tauopathies. Seed amplification assays have demonstrated that these amyloid fibrils are also present in CSF and serum, already before onset of symptoms ^63–67^. In the CSF, their length correlates to disease progression of AD ^60,61^. Monitoring this length may provide an accessible parameter of evaluating disease progression. Especially with current approved therapies targeting amyloid fibrils, they may constitute a relevant readout of therapy efficacy ^17,68–72^.

## Conclusions

FibrilPaint1 is a fluorescein-derivatised peptide that recognises and fluorescently labels Tau amyloid fibrils, but not their monomeric precursors. In combination with Flow Induced Dispersion Analysis (FIDA), a microfluidics technology, we determined the molecular size of amyloid fibrils with sub-microliter sample volumes, including *ex vivo* samples derived from Alzheimer’s Disease, Frontotemporal Dementia and Corticobasal Degeneration patients. This set-up acts as a molecular ruler at layer resolution - we determined Tau fibril length from 4 to 1100 layers in solution. This is an interesting parameter that can be used for diagnostic applications and biochemical research in dementia.

## Methods

### Expression and purification of TauRD

We produced N-terminally FLAG-tagged (DYKDDDDK) human TauRD (Q244-E372, with pro-aggregation mutation ΔK280) in *E. Coli* BL21 Rosetta 2 (Novagen), with an additional removable N-terminal His_6_-Smt-tag (MGHHHHHHGSDSEVNQEAKPEVKPEVKPETHINLKVSDGSSEIFFKIKKTTPL RRLMEAFAKRQGKEMDSLRFLYDGIRIQADQTPEDLDMEDNDIIEAHREQIGG). Expression was induced at OD_600_ 0.8 by addition of 0.15 mM IPTG and incubation at 18 °C overnight. Cells were harvested by centrifugation, resuspended in 25 mM HEPES-KOH pH=8.5, 50 mM KCl, flash frozen in liquid nitrogen, and kept at −80 °C until further usage. Pellets were thawed at 37 °C, followed by the addition of ½ tablet/50 ml EDTA-free Protease Inhibitor and 5 mM β-mercaptoethanol. Cells were disrupted using an EmulsiFlex-C5 cell disruptor, and lysate was cleared by centrifugation. Supernatant was filtered using a 0.22 μm polypropylene filtered and purified with an ÄKTA purifier chromatography System. Sample was loaded into a POROS 20MC affinity purification column with 50 mM HEPES-KOH pH 8.5, 50 mM KCl, eluted with a linear gradient 0-100%, 5 CV of 0.5 M imidazole. Fractions of interest were collected, concentrated to 2.5 ml using a buffer concentration column (vivaspin, MWCO 10 kDa), and desalted using PD-10 desalting column to HEPES pH 8.5, ½ tablet/50 ml Complete protease inhibitor, 5 mM β-mercaptoethanol. The His_6_-Smt-tag was removed by treating the sample with Ulp1, 4 °C, shaking, overnight. The next day, sample was loaded into POROS 20HS column with HEPES pH 8.5, eluted with 0-100% linear gradient, 12 CV of 1M KCl. Fractions of interest were collected and loaded into a Superdex 26/60, 200 pg size exclusion column with 25 mM HEPES-KOH pH 7.5, 75 mM NaCl, 75 mM KCl. Fractions of interest were concentrated using a concentrator (vivaspin, MWCO 5 kDa) to desired concentration. Protein concentration was measured using a NanoDrop™ One^C^ UV/Vis spectrophotometer and purity was assessed by SDS-PAGE. Protein was aliquoted and stored at −80 °C.

### Expression and purification of *Photinus pyralis* Firefly luciferase

We produced His_6_-tag firefly luciferase was expressed in XL10 Gold strain (Stratagene, US). After growing cells at 37 °C, Expression was induced when OD_600_ 0.5, the temperature was lowered to 20 °C. After 45 min incubating, expression was induced by adding 0.10 M IPTG and incubated overnight. Cells were harvested by centrifugation, and pellets were resuspended in lysis buffer (50 mM Na_x_H_y_PO_4_ pH 8.0, 300 mM NaCl, 10 mM β-mercaptoethanol, protease inhibitors (cOmplete, EDTA free, Roche) DNase10 mg/ml). Cells were disrupted using an EmulsiFlex-C5 cell disruptor and lysate was cleared by centrifugation. Super natant was filtered using a 0.22 μm polypropylene filtered and purified with an ÄKTA purifier chromatography System. Sample was loaded onto a POROS 20MC affinity purification column with lysis buffer, and eluted with a linear gradient 0-100%, 5 CV of 0.5 M imidazole. Fractions of interest were collected and dialyzed overnight using dialysis buffer (50 mM Na_x_H_y_PO_4_ pH 8.0, 300 mM NaCl and 10 mM β-mercaptoethanol, 10% glycerol). Protein concentration was measured using a NanoDrop™ One^C^ UV/Vis spectrophotometer and purity was assessed by SDS-PAGE. Protein was aliquoted and stored at −80 °C.

### Peptide synthesis and purification

The peptides were synthesized using a Liberty Blue Microwave-Assisted Peptide Synthesizer (CEM) with standard Fmoc chemistry and Oxyma/DIC as coupling reagents. The peptide concentrations were measured by UV spectroscopy. The peptides were labeled with 5(6)-carboxyfluorescein at their N’ termini. The peptides were cleaved from the resin with a mixture of 95% (v/v) trifluoroacetic acid (TFA), 2.5% (v/v) triisopropylsilane (TIS), 2.5% (v/v) triple distilled water (TDW) agitating vigorously for 3 hours at room temperature. The volume was decreased by N_2_ flux and the peptides precipitated by addition of 4 volumes of diethylether at −20 °C. The peptides were sedimented at −20 °C for 30 minutes, then centrifuged and the diethylether discarded. The peptides were washed three times with diethylether and dried by gentle N_2_ flux. The solid was dissolved in 1:2 volume ratio of acetonitrile (ACN):TDW, frozen in liquid Nitrogen and lyophilized. The peptides were purified on a WATERS HPLC using a reverse-phase C18 preparative column with a gradient of ACN/TDW. The identity and purity of the peptides was verified by ESI mass spectrometry and Merck Hitachi analytical HPLC using a reverse-phase C8 analytical column.

### ThioflavinT aggregation assay

Aggregation of 20 µM TauRD in 25 mM HEPES-KOH pH 7.4, 75 mM KCl, 75 mM NaCl, ½ tablet/50 ml Protease Inhibitor, was induced by the addition 5 µM of heparin low molecular weight in presence of 45 µM ThioflavinT. Impact of peptides was assessed by adding 0.02, 0.2 or 2 µM of peptide. Fluorescent spectra was recorder every 5 minutes, at 37 °C and 600 rpm during 24 hours in CLARIOstar® Plus.

### Fibril extraction

PHFs and SFs were extracted from grey matter of prefrontal cortex from one patient diagnosed with AD according to established protocols ^46^. Tissue was homogenized using a Polytron(PT 2500E, Kinematica AG) on max speed in 20 % (w/v) A68 buffer, consisting of 20 mM TRIS-HCl pH 7.4, 10 mM EDTA, 1.6 M NaCl, 10% sucrose, 1 tablet/10 ml Pierce protease inhibitor, 1 tablet/10 ml phosphatase inhibitor. The homogenized sample was spun from 20 minutes, at 14000 rpm at 4 °C. Supernatant was collected, and the pellet was homogenized in 10% (w/v) A68 buffer. The homogenized was spun once more. The supernatants of both centrifugations were combined and supplied with 10% w/v Sarkosyl, and incubated for 1 hour on a rocker at room temperature. The sample was ultracentrifuged for 1 hour at 100,000 g and 4 °C. The supernatant was discarded, and the pellet was incubated overnight at 4 °C in 20 μl/0.2 g starting material of 50 mM TRIS-HCl pH 7.4. The next day, the pellet was diluted up to 1 ml of A68 buffer, and resuspended. To get rid of contamination, the sample was spun for 30 minutes at 14000 rpm, 4 °C. Supernatant was collected and spun once more for 1 hour at 100000xg, 4 °C. The pellet was resuspended in 30 μl 25 mM HEPES-KOH pH 7.4, 75 mM KCl, 75 mM NaCl, and stored at 4 °C up to a month. Presence of PHF and SF was assessed by TEM.

Narrow filaments were extracted from the grey matter of the middle frontal gyrus from one patient diagnosed with FTD (nhb 2017-019), as described in established protocols ^47^. Fibrils were extracted following the protocol for AD fibrils. After the first ultracentrifugation step, the pellet was resuspended in 250 μl/1g of starting material of 50 mM Tris pH 7.5, 150 mM NaCl, 0.02% amphipol A8-35. The sample was centrifuged for 30 minutes at 3000 g and 4 °C. Pellet was discarded, and the supernatant was ultracentrifuged for 1 hour at 100000xg and 4 °C. The pellet was resuspended in 30 μl of 50 mM TRIS-HCl pH 7.4, 150 mM NaCl, and stored at 4 °C up to a month. Presence of narrow filaments was assessed by TEM.

CBD fibrils were extracted from the grey matter of the superior parietal gyrus of one patient diagnosed with CBD (nhb 2018-007), following established protocols^51^. Tissue was homogenized using a Polytron (PT 2500E, Kinematica AG) on max speed in 20 % w/v 10 mM TRIS-HCl pH 7.5, 1 mM EGTA, 0.8 M NaCl, 10% sucrose. The homogenate was supplied with 2% w/v of sarkosyl and incubated for 20 minutes at 37 °C. The sample was centrifuged for 10 minutes at 20000 g, and 25 °C. The supernatant was ultracentrifuged for 20 minutes at 100000 g and 25 °C. The pellet was resuspended in 750 μl/1g starting material of 10 mM TRIS-HCl pH 7.5, 1 mM EGTA, 0.8 M NaCl, 10% sucrose, and centrifuged at 9800xg for 20 minutes. The supernatant was ultracentrifuged for 1 hour at 100,000 g. The pellet was resuspended in 25 μl/g starting material of 20 mM TRIS-HCl pH 7.4, 100 mM NaCl, and stored at 4 °C up to a month. Presence of CBD fibrils was assessed by TEM.

Brain material of FTD and CBD were obtained from the Dutch Brain Bank, project number 1369. Brain material for AD was donated by prof. J. M. Hoozemans from the VU Medical Centra Amsterdam.

### Electron Microscopy

Fibril samples were diluted to concentrations between 2 µM and 20 µM, and stored in an ice bath until preparation of grids at room temperature. Copper grids with continuous carbon were glow discharged in air of 0.1 bar for 15 sec, using a current of 10 mA, before applying a sample volume of 2.0 µL that was allowed to incubate on the carbon for 60 sec. After incubation, the grid was blotted and subsequently washed twice in 4 µL of milli-Q water and stained twice in 4 µL of uranyl acetate (2w/v% in water), making sure to blot it dry before proceeding to the next droplet. The second droplet of uranyl acetate was left to incubate for 60 sec on the grid, after which it was blotted nearly entirely and left to evaporate for 5 minutes. Imaging took place on a Talos L120C transmission electron microscope from Thermo Fisher Scientific, operating at 120 kV in micro-probe mode. Using a defocus between −1.0 µm and −2.0 µm, the CETA camera recorded images at magnifications of 210, 1.250, 11.000, 36.000, 57.000, and 120.000x, corresponding to an approximate dose between 1 e/ Å² and 50 e/ Å² depending on the exact magnification.

### Size measurement with Flow Induced Dispersion Analysis (FIDA)

Flow Induced Dispersion Analysis (FIDA) was used to determine fibril size. The FIDA experiments were performed using a FIDA1 with a 480 nm excitation source.

Different timepoints preformed TauRD fibrils were diluted to a final concentration of 2 µM in 25 mM HEPES-KOH pH 7.5, 75 mM KCl, 75 mM NaCl, 0.5% pluronic, together with 200 nM of FibrilPaint1. For patient derived Tau filaments, AD, CBD or FTD fibrils were diluted to a final concentration of 2 µM in 20 mM TRIS-HCl pH 7.4, 100 mM NaCl, 0.5 % pluronic (for AD and CBD), or 50 mM TRIS-HCl pH 7.4, 150 mM NaCl, 0.5 % pluronic (for FTD). The mode of operation use is capillary dissociation (Capdis). In this, the capillary is equilibrated with buffer followed by injection of the sample. Only the indicator sample contains fibrils, in order to minimize stickiness to the capillary.

**Table 3.**
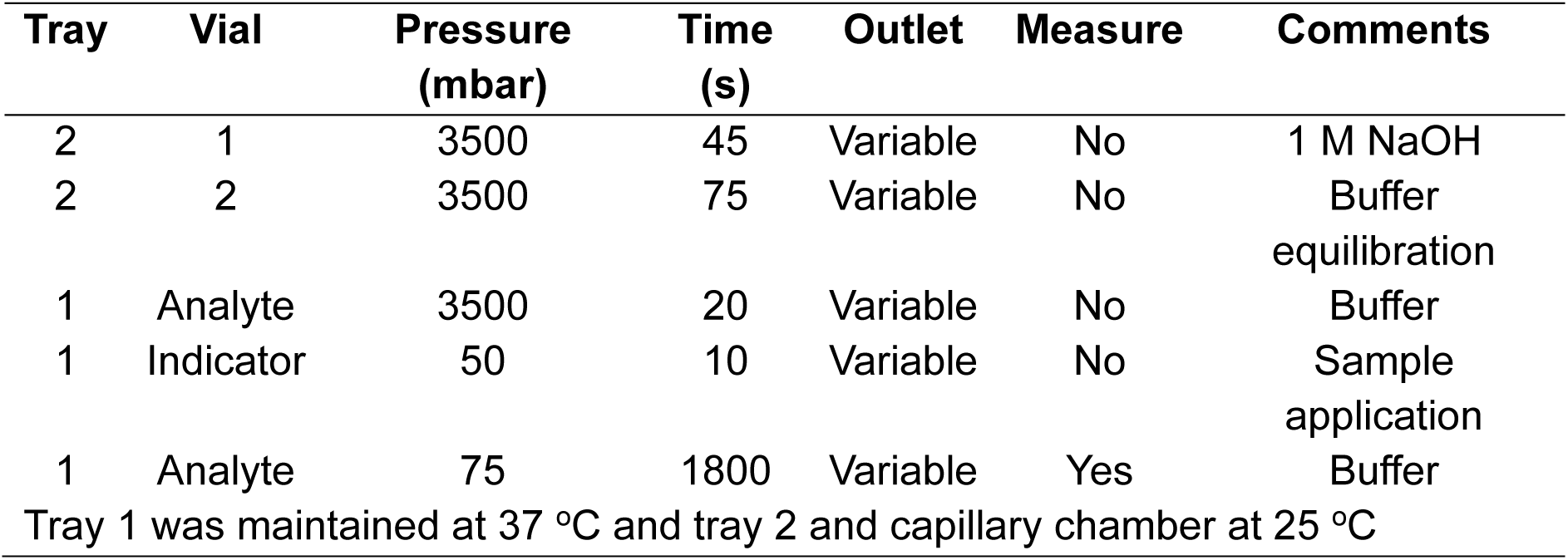
Experimental parameters for Capdis analysis.

### Microscale thermophoresis binding analysis

Binding between FibrilPaint1 and TauRD monomers and fibrils was analyzed using Monolith (NanoTemper Technologies) Microscale thermophoresis system (MST). Thermophoresis was monitored at a concentration of 50 nM of FibrilPaint1 and a dilution series of the sample of interest, in 25 mM HEPES-KOH pH 7.5, 75 mM KCl, 75 mM NaCl. Samples were transferred to premium capillaries (NanoTemper Technologies) and measurements were performed at 37 °C, with medium blue LED power and at MST infrared laser power of 50% to induce thermophoretic motion. The infrared laser was switched on 1 s after the start of the measurement for a 20 s period. Datapoints used were the MST response at 5 seconds.

### Calculation of fibril length from R_h_

For a perfectly spherical particle, the R_h_ is identical to the radius of the sphere. For fibrils, the shape can be approximated as a cylinder. When the length is at least five times as big as the radius (L/R>5), we can apply the Equation 1 ^42,73^:

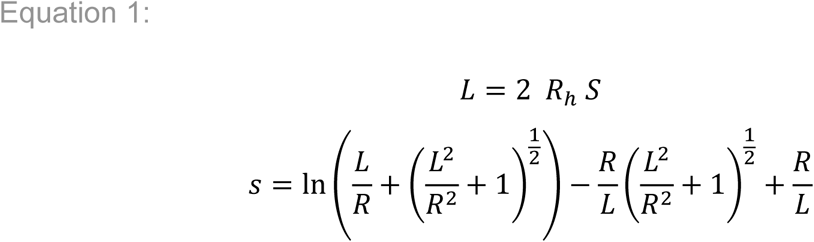

Using equation 1, where L is the length and R the radius of the analysed particle, the mean length of the fibrils as function of R_h_ can be calculated.

As a model, PHF Fibril layers were stacked on top of each other using PyMOL software. 10 structures were created with layers from 1 to 1000 layers. Resulting structures were analysed for their R_h_ using FIDAbio hydrodynamic radii predictor.

The R_h_ were plotted against their layers and analysed with the two-phase association model of GraphPad Prism software v10.4. The resulting mathematical equation can be used to convert the R_h_ to fibril layers, or the R_h_ can therefore be converted to fibril length, as one layer of fibril is 0.47 nm long.

## Acknowledgements

SGDR was supported by grants of the Campaign Team Huntington and Alzheimer Nederland (No. WE.03-2019-03) and a ZonMW TOP grant (No. 91215084). SGDR and FF are principal investigators of the Gravitation Consortium “FLOW” (024.006.036), funded by the the Dutch Ministry of Education, Culture, and Science (OCW). AF thanks The Minerva Center for Bio-Hybrid complex systems and the Saerree K. and Louis P. Fiedler Chair in Chemistry. Measurements on the FIDA1, CLARIOstar® Plus and Monolith were done at the Protein Research Centre of Utrecht University.

## Author contributions

Conception JAP, FAD, TG, GM, AF, SGDR

Design of the work JAP, FAD, SGDR

Acquisition JAP, FAD, MBK, MB, JJMH

Analysis JAP, FAD, HJ, AF, SGDR

Interpretation of data JAP, FAD, HJ, AF, SGDR

Writing original draft JAP, FAD

Revision JAP, FAD, MBK, HJ, TG, JJHM, FF, AF and SGDR

